# AmberMeta: From AMBER Simulation Files to Checked Protocols and FAIR Metadata

**DOI:** 10.64898/2026.11.16.767042

**Authors:** Michele Bonus, Rocco Gentile, Alexander Schug, Holger Gohlke

## Abstract

Molecular dynamics (MD) projects with AMBER often consist of hundreds of runs, and their history is recorded only in the files of these runs. Errors such as a run restarted from the wrong file, a missing or unfinished segment, or a replica that stopped early can therefore pass unnoticed into analyses, Methods sections, and data repositories. Here, we developed AmberMeta, a Python tool that reconstructs the simulation protocol of a finished AMBER project from its files as a draft for the user to review. AmberMeta then checks the reviewed history against the AMBER run outputs and writes a machine-readable record of the runs. This record can serve as findable, accessible, interoperable, and reusable (FAIR) metadata of the simulations, for example for databases such as MDDB or for annotation with ontologies such as MOLSIM, and as the factual basis of a Methods section. In 19 in-house projects, some of them still in progress, AmberMeta reported unfinished runs, replicas at different stages of completion, and runs whose files were not collected with them. In copies of eight projects, it reported 146 of 152 deliberately introduced errors. AmberMeta thus helps improve the reliability, reproducibility, and FAIR data management of AMBER simulations by providing a systematic, file-based record of what was actually simulated.

## Introduction

Simulation results can only be evaluated and reused if it is known how the simulations were run. This matters to reviewers of manuscripts, researchers who build on published simulations, colleagues who take over a project, and authors who return to their own projects years later. The FAIR principles state that research data should be findable, accessible, interoperable, and reusable and should be described with rich metadata and detailed provenance.^1^ The Molecular Dynamics (MD) community has called for simulation data to be shared with such metadata.^2–4^ Databases such as MoDEL, GPCRmd, the NMRlipids Databank, MDRepo, and the federated Molecular Dynamics Data Bank (MDDB) with nodes such as DynaRepo have been created to store simulation data and metadata.^4–9^ Reporting recommendations ask for the simulation software and its version, the size and composition of the system, the thermostat and barostat, the nonbonded cutoff, and how the simulations were split into equilibration and production.^10^ In practice, Methods sections often lack the detail needed to repeat a protocol,^2^ and descriptions of MD data in generalist repositories range from a few words to lengthy free text accounts.^11^ As Musen argued, data-sharing mandates are pointless without appropriate metadata.^12^

With AMBER,^13,14^ one of the most widely used packages for biomolecular simulations, this information is distributed across several files. The topology file (prmtop) describes the system, the control file (mdin) defines a run by specifying the settings, and the output file (mdout) records how the run proceeded, including the used default settings.^15^ A project, defined as the files of one simulated (bio)molecular system in one study, also rarely consists of a single run. Minimization, heating, and equilibration typically precede production.^16^ Production is often split into segments to stay within the job wall-time at computing centers or group clusters,^11^ and each subsegment continues from the restart file written by the predecessor. In addition, each system should be simulated in several independent replicas.^10,17,18^ A project with five replicas of 200 segments each consists of more than a thousand runs.

In such projects, errors can arise from everyday operations. A job script is resubmitted after a crash and reads an older restart file, a restart file is overwritten, a job is killed at the wall-time limit, or a directory is copied to set up a new replica. Each of these errors changes what was simulated. A segment that started from an older restart file covers an overlapping interval of simulation time. A segment missing from the collected files leaves a gap in the trajectory: averages then exclude that segment, and time-dependent analyses must account for the gap. An unfinished segment shortens the simulation and leaves replicas with different amounts of sampling. ^19^ None of these errors shows in the control files: the mdin of a segment states how many steps to run and that the run continues a previous one, but not which restart file it reads, which is given on the command line, nor the time at which it starts, which AMBER takes from that restart file.^20^ Documentation written by hand thus easily drifts from what was run.^21^ Whether the runs of a finished project form the protocol that its authors report can therefore only be checked run by run, which is rarely feasible for hundreds of runs.

Today, authors typically compile such information manually for their Methods sections, and depositors do the same for repository forms; the input file of the MDDB workflow, for example, asks the depositor for the program and its version, the frame interval, the time step, the temperature, the ensemble, the force field, and the water model.^22^ Tools that extract metadata from simulation files exist, but to our knowledge, no published tool checks the runs of a finished AMBER project for internal consistency against the output files that AMBER wrote for these runs (Table 1). GROMACS MetaDump derives metadata from the input file of a single GROMACS run and validates them against a JSON Schema.^23^ MDverse indexes MD data deposited in public repositories, but its published analysis parsed only GROMACS files.^11^ For AMBER, iBIOMES Lite summarized a simulation directory as a tree of minimization, heating, equilibration, and production in a fixed order,^15^ and the IDA database extracts metadata from uploaded AMBER topology, trajectory, and output files, checks that they are complete, and is designed to check that their atom and frame counts agree.^24^ Of the AmberTools programs, cpptraj reads topology and trajectory files and parmed edits topology files, but the user orders and joins the runs.^25,26^ Workflow systems such as AiiDA record provenance, and the BioExcel Building Blocks provide an interface for it, but only for simulations run through them.^27,28^

**Table 1:** Tools That Describe the Runs of AMBER or GROMACS Simulations from Their Files.

|  | iBIOMES Lite<sup>15</sup> | GROMACS MetaDump<sup>23</sup> | IDA<sup>24</sup> | AmberMeta (this work) |
|---|---|---|---|---|
| MD engines | AMBER, GROMACS, NAMD<sup>a</sup> | GROMACS | AMBER | AMBER |
| input | simulation directory | run input file (.tpr), optionally topology and structure | uploaded topology, trajectory, and output files | simulation directory |
| runs of a project | tree of minimization, heating, equilibration, and production processes in fixed order | single run | one simulation record; replica exchange runs split between exchanges | phases and runs in any order, each with the file it started from |
| replicas | replica exchange runs grouped by folder or naming pattern; one subfolder per copy | not described | replica exchange, thermodynamic integration, and umbrella sampling windows | independent replicas, inferred from the directory layout or declared |
| checks | unfinished tasks flagged; checks between runs not described | metadata validated against a JSON Schema | file completeness; force field against its database; atom and frame counts planned | input file recorded by AMBER, completion, mdin against mdout, continuity in time, gaps in numbered series, consistency between replicas |
| metadata format | XML and HTML summaries | JSON or YAML with a JSON Schema | database records; XML metadata following DataCite 4.5; persistent identifiers planned | manifest and summary in JSON or YAML, methods summary in JSON and Markdown; no public JSON Schema |
| interface | command-line scripts; static HTML | web application, API, command line | web database | command line; local, browser-based editor |

<sup>a</sup> Also the quantum chemistry codes Gaussian and NWChem.

We developed AmberMeta to help researchers determine whether the available run files support the reported history of each replica in an AMBER project (Figure 1). Its discover command drafts the protocol from the files, and the user reviews the draft in AmberMeta’s editor and modifies it where needed. The validate command then checks the protocol against the output files that AMBER wrote for every run, and the plan command writes a record of the runs. AmberMeta is written for the researchers who ran the simulations and for the principal investigators and data stewards who archive them. It has three main intended uses: (i) to perform a sanity check of a project when a simulation campaign ends, before its results are analyzed and published and before its data are archived; (ii) to extract metadata on the simulation part of a dataset for repositories, for databases such as MDDB, and for ontologies such as MOLSIM; and (iii) to provide the factual basis of a Methods section.

**Figure 1:**
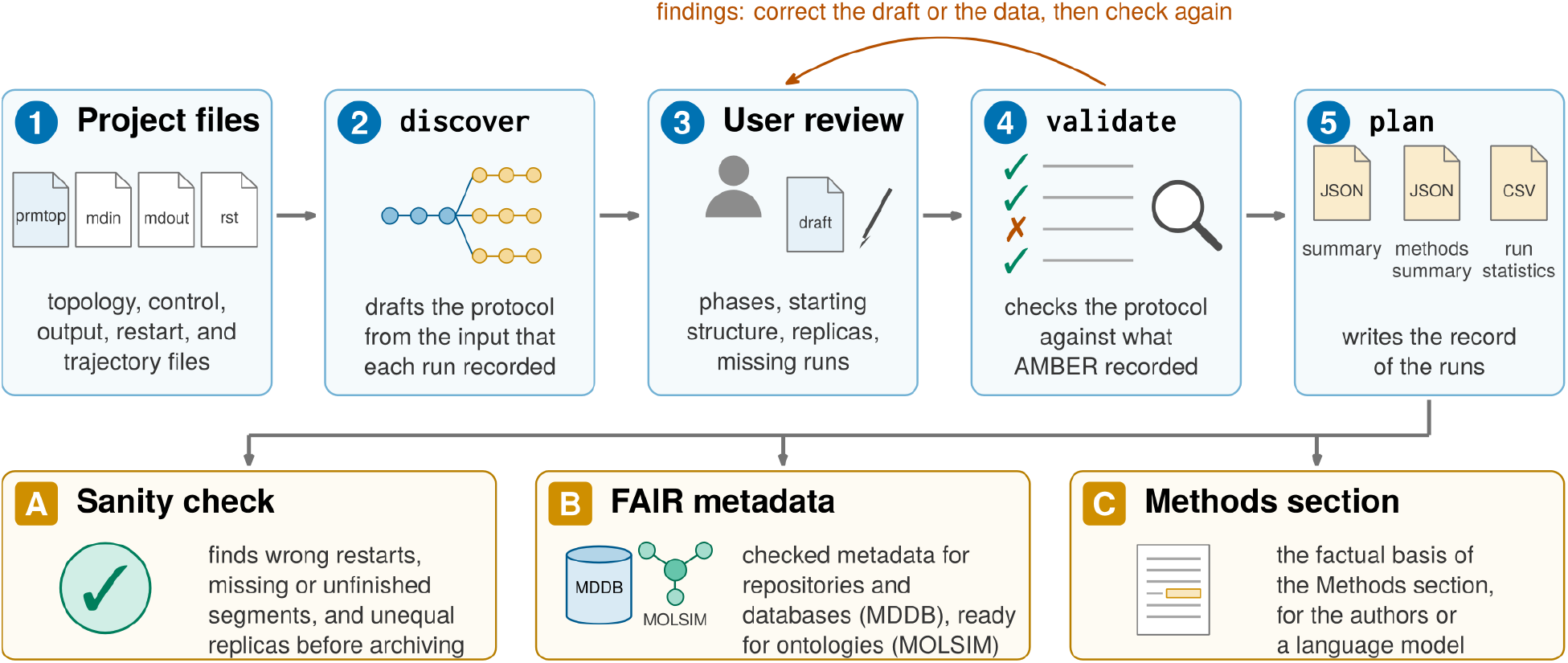
AmberMeta’s workflow and its uses. (1) The files of a finished AMBER project. (2) discover drafts the protocol as a *manifest* and links each run to the input that its mdout records. (3) The user reviews and corrects the draft. (4) validate checks the protocol against the output files of the runs and reports *findings*, which lead back to the review. (5) plan writes the record of the runs: the *summary*, the *methods summary*, and a table of run statistics. The record serves (A) as a sanity check before analysis and archiving, (B) as FAIR metadata for repositories, databases, and ontologies, and (C) as the factual basis of a Methods section. The numbers correspond to the subsections of How AmberMeta Works and the letters to those of the Results.

We demonstrate AmberMeta on a sample project distributed with the software and apply it to 19 projects of an in-house test collection, among them the published simulations of the taurine transporter TauT.^29^ We also test whether AmberMeta reports errors deliberately introduced into copies of eight of these projects and whether language models can draft a Methods section from its record without adding facts. AmberMeta can be installed with pip install ambermeta and is available at https://github.com/CPCLab/ambermeta.

## How AmberMeta Works

AmberMeta works in five steps (Figure 1). The user points it at the directory of a project (step 1), discover drafts the protocol (step 2), the user reviews the draft (step 3), validate checks the protocol against the output files of the runs (step 4), and plan writes the record (step 5). The three uses described in the Introduction all build on this record. The terms that AmberMeta uses for the parts of a protocol are defined in Table 2 and set in italics throughout. The rules behind each step are given in the Supporting Information (Sections S1 to S4).

**Table 2:** Terms Used by AmberMeta and in This Article^a^.

| term | meaning |
|---|---|
| project | the files of one simulated (bio)molecular system in one study, usually in one directory tree |
| *Simulation* | AmberMeta’s description of a project: its topologies, its starting structure, and its runs |
| role | the stage of the protocol to which a run belongs: minimization, heating, equilibration, or production |
| *Phase* | consecutive runs with the same role |
| *Step* | one run, that is, one execution of pmemd or sander with one mdin and one mdout |
| segment | a production run that continues the previous run from its restart file |
| replica | an independent copy of a simulation, usually continuing from shared preparation runs |
| *manifest* | the text file (JSON or YAML) that lists the *Phases* and *Steps* of a *Simulation* with their files, the file from which each run started, and the replicas |
| *finding* | an issue that validate reports on a run, a replica, or the project |
| record | the files that plan writes: the *summary*, the *methods summary*, and a table of run statistics |
| discover,<br>validate,<br>plan | the commands that draft, check, and record the protocol |

<sup>a</sup> Terms in italics are defined in the section “Drafting the Protocol”.

### Drafting the Protocol (Steps 1 to 3)

Writing down the protocol of a project with hundreds of runs by hand is tedious and error-prone, so AmberMeta drafts it. It describes a protocol with three nested terms that follow the way a Methods section describes simulations (Table 2): the simulated system (*Simulation*), the stages of the protocol (*Phase*), and the single runs (*Step*). A *Simulation* holds the topologies of a project and its starting structure. The starting structure is the coordinate file from which the first run started, usually the output of the system preparation. Each topology is labeled as standard or as a hydrogen mass repartitioning (HMR) topology.^30^ A *Simulation* can thus hold a standard topology for equilibration with a time step of 2 fs and an HMR topology for production with 4 fs. Each run has a role, the stage of the protocol to which it belongs: minimization, heating, equilibration, or production. Besides raising the temperature, the heating stage can also adapt other settings, for example, switch on pressure coupling. As is common in the field, we use the term equilibration for a stage of the protocol, without implying that the system is equilibrated afterward. AmberMeta keeps heating apart from equilibration, although some authors count heating as part of the equilibration, and the user can change the role of any run. Consecutive runs with the same role form a *Phase*, and each run is a *Step*. A *Step* names its topology, its mdin and mdout, its trajectory and restart file, and its input coordinates: the starting structure, the restart file of another *Step*, or another file. When one *Step* names another’s restart file as its input, the protocol links the two runs. These links form a graph that can represent shared preparation runs and highlight where replicas branch from a joint restart file (Figure 2A). The *manifest* stores this graph as a JSON or YAML file that people and programs can read and that any text editor can change.

**Figure 2:**
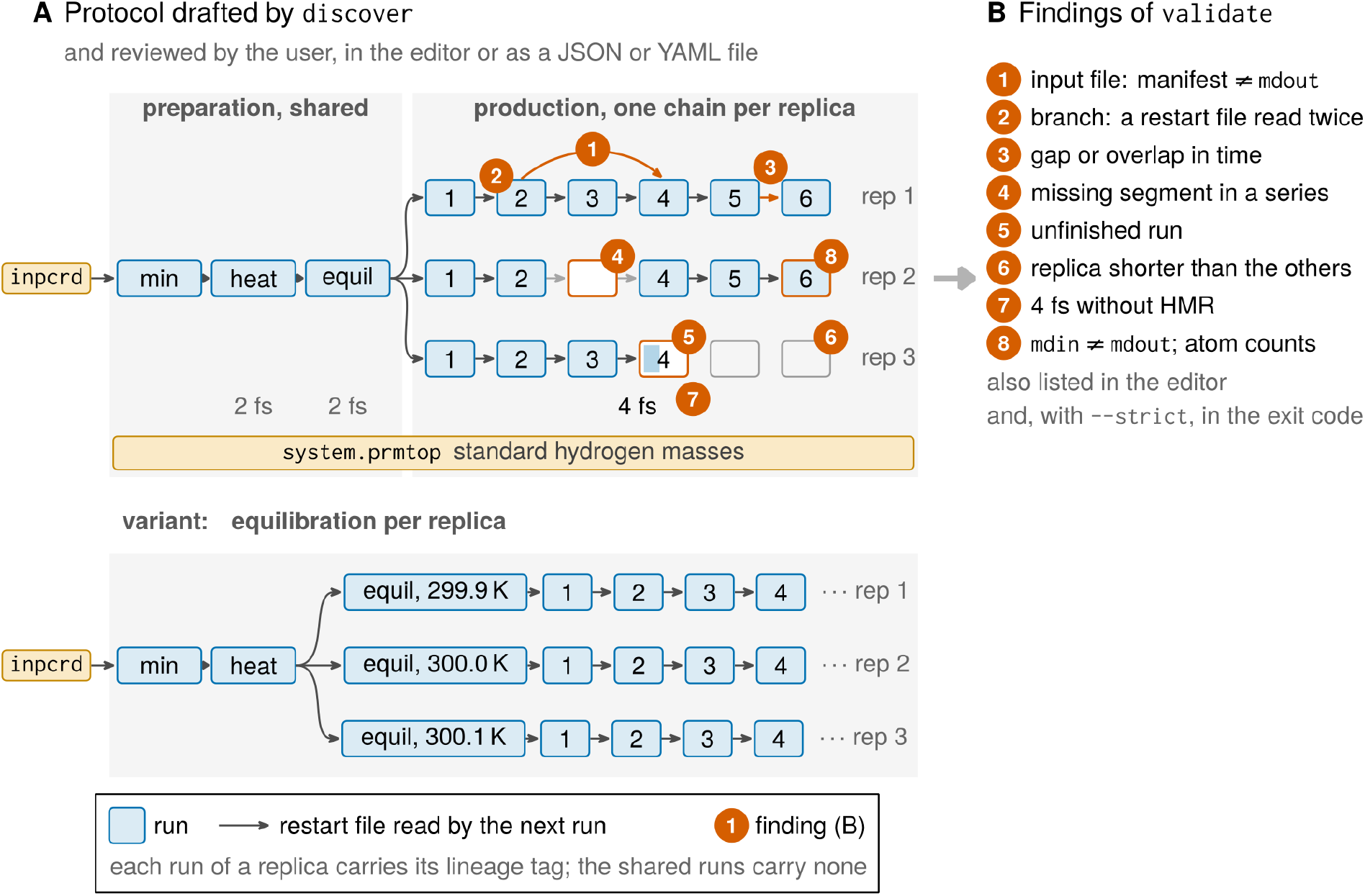
The protocol of a project and the *findings* of validate, shown on an exemplary project. (A) The protocol as a graph of runs linked by restart files. The minimization, heating, and equilibration runs are shared, and three replicas continue from the restart file of the shared equilibration as chains of production segments. Below, a variant in which each replica has its own equilibration, here at slightly different target temperatures. Orange numbers mark where the *findings* of panel B apply. *Findings* 1 and 2 show how validate reports one error, depending on the *manifest*. In replica 1, segment 4 read the restart file of segment 2, as its mdout records (dashed orange arrow), and not that of segment 3. If the *manifest* declares the restart file of segment 3 (solid arrow), validate reports *finding* 1 on segment 4. If the *manifest* follows the mdout, as the draft of discover does, the dashed arrow replaces the solid one. Segments 3 and 4 then both continue from segment 2, and validate reports this branch as *finding* 2. (B) The *findings*: (1) the *manifest* declares another input file than the mdout records, (2) two runs read the same restart file, (3) a gap or overlap in simulation time, (4) a segment missing from a numbered series, (5) an mdout without the final timing summary, (6) a replica shorter than the others, (7) a time step of 4 fs on a topology with standard hydrogen masses, and (8) mdin settings that differ from those echoed in the mdout, or atom counts that differ between files.

Each type of file contributes part of the description of a run; the AMBER manual defines the file formats and settings.^20^ From the topology, AmberMeta reads the atom count, the residue composition including water molecules and ions, the net charge, and the hydrogen masses, from which it labels the topology as standard or HMR. From the mdin, it reads the settings of the run and derives from them the ensemble, thermostat, barostat, cutoff, bond constraints, and positional restraints. It also flags methods that need separate reporting, such as replica exchange, Gaussian accelerated MD, free energy calculations, and constant pH. From the mdout, it reads the program and its version, the graphics processing unit (GPU), whether the run finished, and the temperature, pressure, density, and energy of every energy record. Restart files and NetCDF trajectories store the simulation time of their coordinates,^20,31^ which AmberMeta uses to check that consecutive runs connect in time. AmberMeta also works on incomplete projects. A run without an mdin is described from the settings that its mdout echoes, a run without an mdout is listed as prepared but not run, and a file that is missing or cannot be opened is noted on its run while AmberMeta continues with the others (Supporting Information, Section S1).

The discover command drafts the *manifest* from the directory of a project. It groups the files of each run by their common name and assigns each run a role, first from the mdin (imin = 1 requests a minimization), then from the words min, heat, equil, or prod and their variants in file and directory names, and then from the settings, for instance positional restraints during equilibration. To link the runs, discover uses the File Assignments block at the top of each mdout, in which AMBER records the input coordinates (INPCRD) that the run actually read, whatever the job script or the file names suggest. discover then links each run to the run that wrote this file, so that the draft follows the order in which the runs were actually executed. Where this record is unusable, for example because AMBER cut a long path short, a run is linked to the run before it in its directory, in the order of the file names, in which prod_2 precedes prod_10. With several topologies, each run is assigned the topology whose atom count matches its mdout. Replicas are recognized from the directory layout: sibling directories that hold the same series of runs (rep1 to rep3, for instance) become replicas, whereas a directory with the preparation runs that all replicas share, such as the minimization and heating, stays outside them. The documentation that comes with the software recommends a layout for new projects that discover recognizes (Supporting Information, Section S2).

The draft cannot replace the user’s review, because directory layouts and file names follow no common rule,^15^ and the stages of a protocol are not always unambiguous.^16^ discover therefore reports the inferences (the starting structure, the role of each run, the replicas, and the indices missing from numbered series) it used to build the draft (marked applied) and the questions it leaves for the user (marked needs_you). The user checks these items and corrects the draft in AmberMeta’s browser-based editor or in a text editor. Replicas in layouts that discover does not resolve are declared in the same way, for example independent simulations at several temperatures or with several ligands, each with its own replicas in subdirectories (300K/rep1 beside 310K/rep1). Where a project is organized so unusually that the draft would need more correction than it saves, the protocol can be built by hand in the editor: the user creates the *Phases* and runs, drops the files into them, and declares the replicas. validate and plan then work as on a draft.

### Checking the Protocol (Step 4)

To find the errors described in the Introduction, the validate command checks the protocol at three levels: each run, consecutive runs, and the replicas of a *Simulation*. Most checks compare the files of the runs with each other, for example the mdin of a run with the settings that its mdout echoes, or the end of one run with the start of the next. One check compares the *manifest* with the mdout files: the input file that the *manifest* declares for a run has to be the one that AMBER recorded in the mdout of that run as read. A draft of discover normally passes this check, because it follows the same record, but a *manifest* that the user edited or wrote by hand need not. Each failed check gives a *finding*, a message attached to the run or replica concerned. Figure 2 lists the kinds of *findings* and shows where in a protocol each of them applies; the Supporting Information describes every check (Section S3).

For each run, AmberMeta compares the input coordinates that the *manifest* declares with those that the mdout records; they differ, for example, when a job script read an older restart file than the protocol states. It reports runs whose mdout lacks the timing summary that AMBER writes at the end of a run, because the run stopped early or is still running. It compares the atom counts of the topology, coordinates, output, and trajectory, which differ when a run used the topology of another system. It also compares the number of integration steps, the time step, and the trajectory write interval of the mdin with those that the mdout echoes, which differ when the mdin was edited after its run. Finally, it reports a time step above 2 fs on a topology with standard hydrogen masses. Because time steps of 4 fs rely on HMR,^30^ such a run either used an unsuitable time step or used an HMR topology other than the one in the *manifest*.

Between consecutive runs, AmberMeta checks that the simulation time continues. For each run that starts from a restart file, it compares the time stored in that file with the end of the run that wrote it. A gap means missing simulation time, and an overlap means that a stretch was simulated twice, for example after a restart file was overwritten by an older one. Differences within a small tolerance are accepted, and gaps that the user declares as intended are not reported. Because the draft follows the recorded input coordinates, a run that read an older restart file stays consistent in time. It shows instead as a branch, two runs of the same replica that continue from the same restart file, and AmberMeta reports both. In a numbered series such as prod_0001 to prod_0200, AmberMeta reports the indices missing between the lowest and the highest index. Across the replicas of a series, it uses their combined range, so that a replica that stopped before the others is reported as well.

Replicas should describe the same system under the same conditions. AmberMeta reports replicas that differ in atom count, and it warns when replicas differ in time step, cutoff, thermostat, or pressure coupling in runs of the same role, or in target temperature during production. Some protocols equilibrate each replica at a slightly different temperature before all continue at a common production temperature (Figure 2A, below); such replicas are not reported. This is not replica exchange, in which replicas at different temperatures exchange configurations during the run; AmberMeta flags replica exchange runs but does not model them. Replicas often start from the same coordinates with random initial velocities,^32^ and two replicas run with the same random seed are not independent. AmberMeta therefore warns when replicas that continue from the same restart file used fewer distinct random seeds than there are runs. For use in scripts, validate ends with a nonzero exit code when files named in the *manifest* are missing or cannot be opened or when atom counts differ between replicas. With the ––strict option, every other *finding* also leads to a nonzero exit code, so that a script can, for example, stop before a project is archived. A *finding* that the user has checked, for example a replica that was planned to be shorter, can be acknowledged with a note. The note is kept in the *manifest* and in the record, and the strict check no longer counts that *finding*. also The editor shows the same *findings* beside the runs (Figure 3).

**Figure 3:**
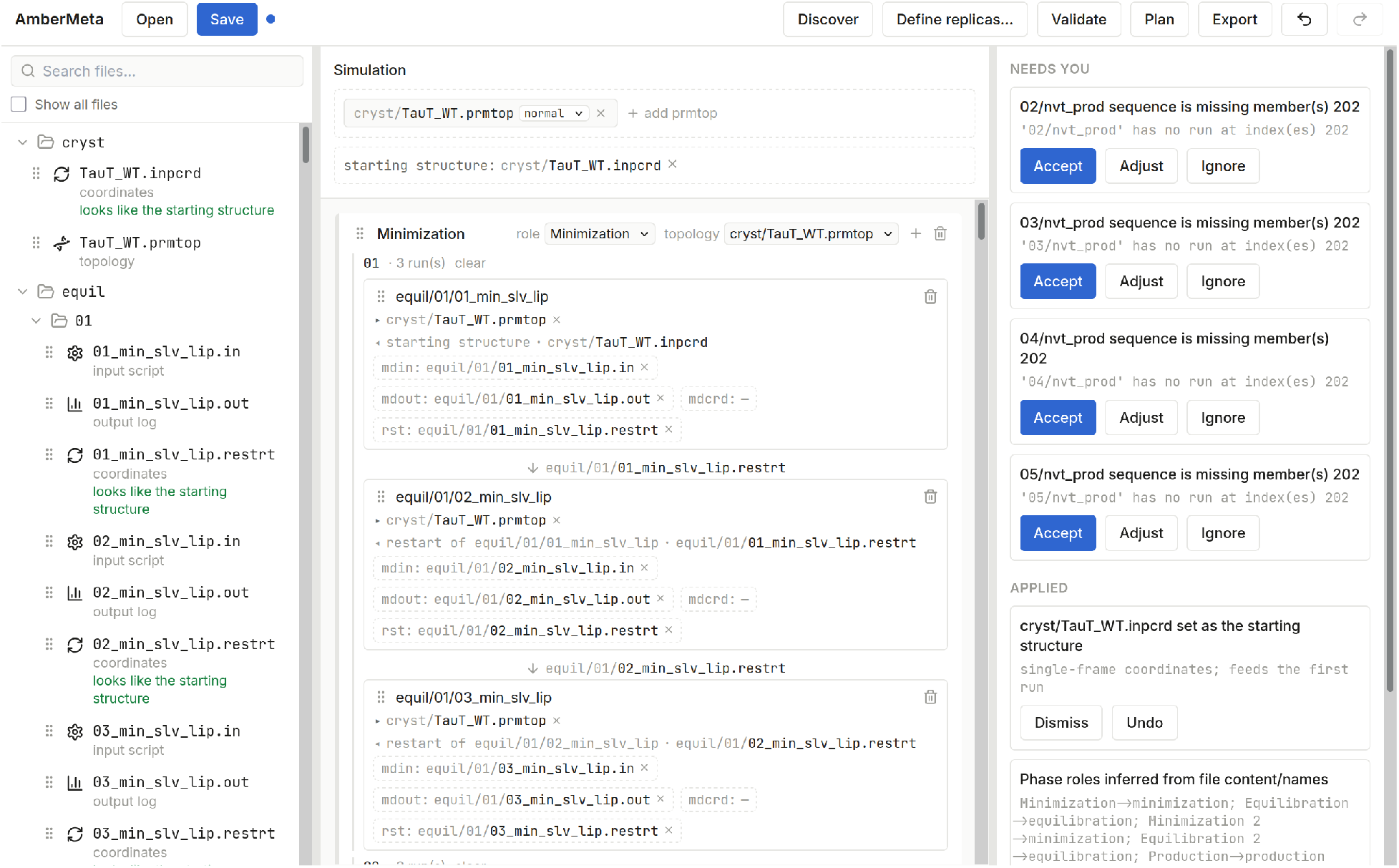
AmberMeta’s editor for an in-house project: simulations of the human taurine transporter TauT in five replicas (01 to 05),^29^ after discover and the acceptance of the proposed replicas. Left, the project files and their types. Center, the topology, the starting structure, and the first runs of the minimization of replica 01, each with the restart file from which it continues. Right, the *findings*, which the user can accept, adjust, or ignore. They list run 202 as missing in replicas 02 to 05, because replica 01 ran one production segment more than the others and queued run 202.

### Writing the Record (Step 5)

To make the checked protocol reusable, the plan command writes its record in three parts, each for one use. The *summary*, a single file in JSON or YAML format, is the full record to be archived with the data: the parsed settings and *findings* of every run, the values that each mdout echoes, and the simulated time per replica, per role, and in total. These times count what the mdout files show was run, not what the mdin files requested, and runs that were prepared but have no output, for example because they are still queued, are listed separately. The *methods summary*, in JSON and as a Markdown file to read, condenses the *summary* into a description of the protocol for the authors of a Methods section and for language models. A table in comma-separated values (CSV) format lists the simulated time of every run with the mean and standard deviation of its temperature, pressure, density, and total energy, for a quick look at every run.

The *methods summary* also describes the system: the residues by class, the water molecules with the number of atoms per water molecule (e.g., three for TIP3P, four for models with an extra point such as OPC), the ions, the box, and the hydrogen masses. It lists the *Phases* in the order in which they ran, with their runs, replicas, and simulated time, and it states each setting once per *Phase*, with the number of runs that share it and the values of the runs that differ, such as a series of decreasing restraint force constants. Each setting carries its source: set in the mdin, echoed in the mdout because the mdin left it at its default, an AMBER default that neither file states, or derived by AmberMeta from other settings, such as the ensemble. The *methods summary* also groups the *findings* and lists what AMBER run files do not contain, among them the force field and the water model. The *methods summary* takes less than 20 kB even for projects whose *summaries* take tens of megabytes. Together, the *summary*, the *methods summary*, and the table of run statistics serve as a sanity check of the project, as input for metadata in databases and ontologies, MDDB and MOLSIM among them, and as the factual basis of a Methods section (Results).

### Installing and Running AmberMeta

AmberMeta is written in Python, requires version 3.9 or newer, and can be directly installed with pip install ambermeta, with the editor as the optional extra ambermeta[gui]. It runs on the user’s computer, from the command line or in a browser-based editor that needs no internet connection, and unpublished data stay on the machine. The command line suits scripts, for example, a check at the end of every simulation campaign. Reading NetCDF files requires the netCDF4 or SciPy package; without them, AmberMeta takes the simulation times from the mdout files and cannot check the restart files themselves; an overwritten restart file then goes unnoticed. A quickstart guide and the documentation of every command come with the software.

## Results

### Sanity Check of a Finished Project

The first use of AmberMeta is a sanity check of a finished project. We tested it on three kinds of data: the sample project distributed with the software, copies of the sample and of in-house projects into which we had introduced known errors, and an in-house test collection as we found it. All results were obtained with AmberMeta 1.3.0; the procedures are given in the Methods, and the values per project, the outputs, and the run times are in the Supporting Information and its data files.

#### The Sample Project

The sample holds five consecutive production segments of 20 ns each of human YKL-40 (chitinase-3-like protein 1) in complex with chitohexaose, a solvated system of about 65,000 atoms, run with the GPU engine of AMBER 22, pmemd.cuda, and a time step of 4 fs.^33–35^ discover reconstructed the protocol as the runs had been executed: one production *Phase* of five segments, each continuing from the restart file of its predecessor and the first from the file that its mdout records. The record gives 100 ns of simulated time, and the mean temperatures and densities of the five segments agree within their standard deviations (Supporting Information, Section S5).

**validate** reported one *finding*: every segment used a time step of 4 fs, but the topology in the sample has standard hydrogen masses. The project had used a standard topology for its runs with 2 fs and an HMR topology for those with 4 fs, and only the first was collected into the sample. The mdout files show that the runs read the HMR topology. AMBER truncates long paths in the mdout, so the name of the topology is incomplete there. However, the first line of a topology file gives the date and time at which the file was written, and the mdout repeats this time stamp for the topology that the run read. In all five mdout files, this time stamp is three minutes later than that of the topology in the sample and equals that of the HMR topology. AmberMeta does not yet compare these time stamps. In a copy of the sample whose topology has raised hydrogen masses, validate reported no *finding*. The case shows a problem that matters for research data management: a dataset can look complete and still lack a file that is needed to reproduce it.

#### Introduced Errors

To test the checks on known errors, we introduced the synthetic errors listed in Table 3, each a typical slip of everyday work, into copies of the sample and of eight in-house projects and checked each copy as a user would. In the sample, every error was reported, and each left its own trace: a missing segment appears as a gap in the numbered series (error 1), a run that read an older restart file as a branch, where two runs continue from the same file (error 2), an overwritten restart file as an overlap in simulation time (error 4), and an interrupted run as unfinished (error 5). AmberMeta also tells whether an error lies in the protocol or in the simulation. When the *manifest* names the wrong restart file (error 3), AmberMeta reports that it differs from the file that the mdout records; when the run itself read the wrong file (error 2), the draft follows the mdout, and the branch is reported. The strict check failed for every modified copy, so that a script would stop before such a project is archived.

**Table 3:** Errors in Multi-Run AMBER Projects, the *Findings* by Which AmberMeta Reports Them, and the Errors Reported When Introduced into Copies of the Sample and of In-House Projects.

|  | error | how it was introduced | *finding* | sample<sup>a</sup> | in-house projects<sup>b</sup> |
|---|---|---|---|---|---|
| 1 | segment missing (not collected, deleted, or never run) | files of one segment deleted | index missing in the numbered series; the next segment recorded an input file that the *manifest* does not name | yes | 20/24 |
| 2 | run read an older restart file (job resubmitted) | mdout of a segment records the restart file of the segment before its predecessor | branch: two runs continue from the same restart file | yes | 24/24 |
| 3 | protocol states the wrong restart file (error in the *manifest*) | *manifest*: a segment reads the restart file of the segment before its predecessor | recorded input differs from the declared one; branch | yes | 23/24 |
| 4 | restart file overwritten by an older one | restart file of a segment replaced by that of its predecessor | overlap in simulation time | yes | 23/24 |
| 5 | run stopped early (wall-time limit, crash) | mdout truncated before its final summary | unfinished run | yes | 24/24 |
| 6 | replica stopped early | one replica holds segments 1 and 2, the other 1 to 5 | indices 3 to 5 missing in the shorter replica | yes | 9/9 |
| 7 | mdin edited after its run | number of integration steps doubled in the mdin | mdin differs from the values echoed in the mdout | n.t. | 20/20 |
| 8 | run used the topology of another system | topology with a different atom count bound to a run | atom counts differ | n.t. | 3/3 |

<sup>a</sup> Copies of the sample; the strict check failed in every case. n.t., not tested. The exact modifications and the messages are given in the Supporting Information (Section S5).
<sup>b</sup> Errors reported on the run that carried them, of errors introduced at randomly chosen runs of copies of eight in-house projects (Supporting Information, Section S6). In total, 146 of 152 errors were reported.

Real projects vary more in layout, file names, and history than a sample prepared for the purpose. In copies of eight in-house projects, AmberMeta reported 146 of 152 introduced errors (Table 3, Figure 4A), among them every edited mdin, every run bound to the topology of another system, and every replica that stopped early. The six misses had three causes, and each points to something the user can do. Three occurred in a project whose replicas have one directory for production at the top level and a differently named one for equilibration inside a common equilibration directory. From this layout, AmberMeta infers no replicas, and a run with the same index in another replica then masked the gap. Declaring the replicas in the editor or the *manifest* prevents this, and the documentation recommends a layout that discover recognizes. Two misses occurred where the next run had been set up but had not yet run. Only a run that has run records which file it read, so nothing showed the error. One concerned a restart file overwritten in a chain of minimizations; minimizations do not advance the simulation time, so there was no time to compare. Further *findings* on neighboring runs were rare and occurred in one project. The misses and these *findings* are listed per project in the Supporting Information (Section S6).

**Figure 4:**
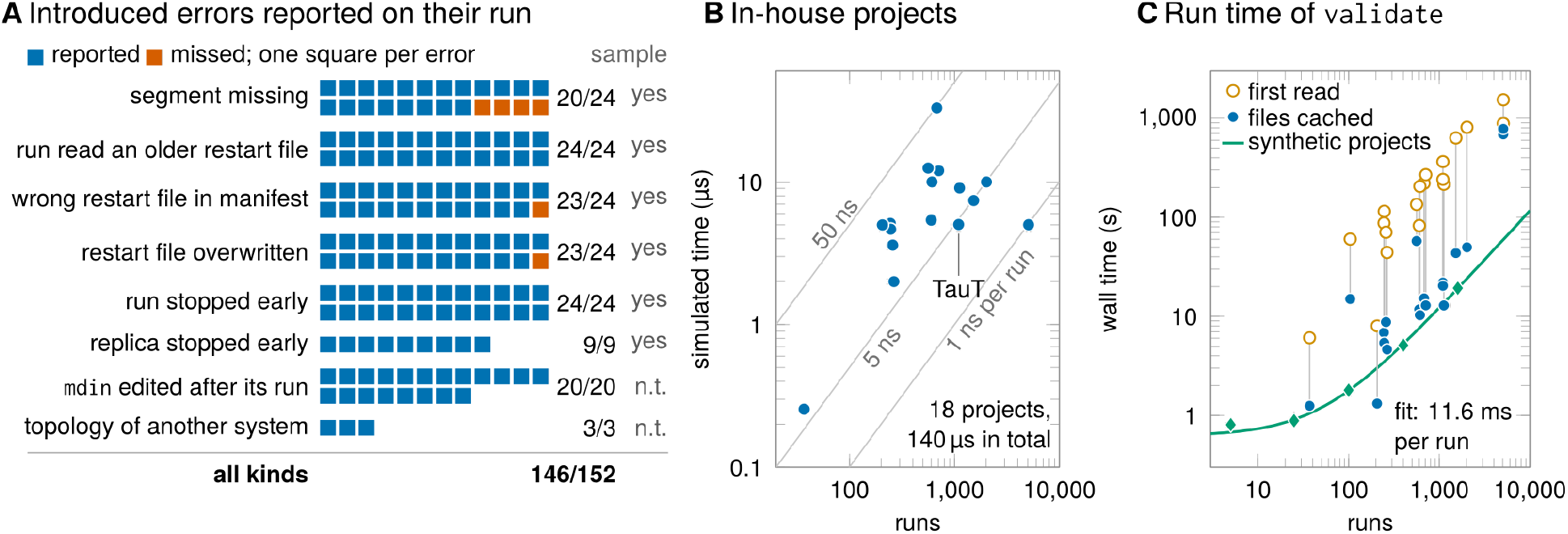
Evaluation on in-house projects. (A) Errors introduced into copies of eight in-house projects, one square per error: blue if AmberMeta reported it on the run that carried it, red if not (146 of 152 in total). The right column gives the result for the same kind of error in a copy of the sample (n.t., not tested); see also Table 3. (B) Runs and simulated time of the 18 in-house projects whose runs have mdout files; gray lines mark a mean simulated time per run of 1, 5, and 50 ns. Two pairs of projects coincide, among them the two TauT projects. (C) Wall time of validate against the number of runs for all 19 in-house projects with runs, when their files were first read from the network file system (open circles) and with the files cached (median of the three repetitions after it, filled circles). Diamonds mark synthetic projects built from the sample, with a least-squares fit of 0.62 s plus 11.6 ms per run (Supporting Information, Section S6).

#### An In-House Test Collection

Most MD data today are never put into a public repository. The data sets behind most publications are typically not directly available.^36^ When trajectories are kept only in local storage after publication, they can gradually become inaccessible as computing systems are replaced, hardware fails, or storage media deteriorate.^4^ Trajectories can also be preserved and managed according to FAIR principles^36^ on storage maintained by research groups or computing centers.^15^ To develop and test AmberMeta and other tools for research data management, members of our laboratory assembled a collection of MD projects run over about the last ten years, published and unpublished. The collection is a small part of our group’s MD data and was meant to represent the states in which MD data are found during and after a study. We applied AmberMeta to it as a data steward or principal investigator would before archiving a project or depositing it in a public repository. Of the 50 project directories that were set up as the framework of the collection, 19 have been filled with simulation runs so far, and one could not be opened. The 19 projects hold more than 100,000 files and 5.3 TB in total, with tens to thousands of runs and up to 15 replicas per project (Figure 4B). AmberMeta read all of them without a failed file and drafted more than 20,000 runs.

validate reported *findings* in 9 of the 19 projects, and they show the states in which MD data are found while a study runs or after it has ended. Replicas had reached different stages: in two projects, one replica had run fewer production segments than the others, in one of them fewer than half. From the files alone, AmberMeta cannot tell whether the shorter replica stopped early or the others ran longer than planned; this is a question for the researchers. Some runs had not finished, because they stopped early or were still running when their files were collected. In one project, the runs used a time step of 4 fs, but the only topology in the collection has standard hydrogen masses; whether the runs used an HMR topology that was not collected cannot be decided from the files. In another, HMR topologies were present, but the production runs had no mdin files, and the draft bound them to a standard topology with the same atom count, a choice that the user corrects in the review. In two other projects, the trajectory of one equilibration run per replica did not cover the length of the run, and the replicas ran production at different target temperatures, which may have been intended. A few runs had read restart files whose producing runs are not in the collection, and in one run, the time step in the mdin differed from the one that its mdout records, so the mdin had presumably been edited after the run. These are the questions that a data steward would ask the researchers before the data are archived or deposited, and the researchers who ran the projects confirmed the *findings*. The *findings* per project are given in the Supporting Information (Section S6).

#### A Published Multi-Stage Project: TauT

The TauT projects show how AmberMeta handles a large published project and why its draft has to be reviewed. They hold the simulations of the human taurine transporter TauT and its variant p.A78E in a membrane, five replicas of each, which have been published.^29^ Each system is one project of more than 5,000 files. For each replica, one directory holds the minimization, heating, and equilibration runs, and a second directory holds about 200 production segments. discover drafted the protocol of each system within seconds and proposed the five replica directories as replicas, which the user confirmed in the editor.

The draft also showed two cases that only the user can resolve. The heating run of each replica, 04_nvt_heat, lies in a directory named equil. Because discover reads directory names before file names, it assigned the run to equilibration, which the user corrects in the editor. Conversely, the production segments are named nvt_prod, but their mdin files request constant pressure, and the record describes them as NPT runs, as they were run. The record also shows that the runs of each system were spread over three GPU models (GeForce GTX 980 Ti, GeForce GTX 1080, and Tesla K20m).

validate reported that the replicas differ in length (Figure 3): replica 01 had run one production segment more than the other four. No data are missing; the replicas differ by one segment of 5 ns.

#### Run Time

A check fits into the end of a simulation campaign. When the files were first read from the network file system, validate took a few minutes for a typical in-house project and less than half an hour for the two largest projects, which hold about 5,000 runs each (Figure 4C). With the files cached, validate took less than a minute for most projects, about an order of magnitude less than at the first read, and discover took seconds. For the two largest projects (130 GB and 139 GB), the files did not stay in the cache, and validate still took more than ten minutes, most of which were spent reading files from the network file system. The time grows linearly with the number of runs, and the memory needed is set by the size of the topology file. The times and memory per project are given in the Supporting Information (Section S6).

### FAIR Metadata

The second use of AmberMeta is to extract metadata on the simulation part of a dataset. Its record holds most of the setup items that a reporting checklist asks for and most of the simulation fields of the MDDB input file (Table 4; item by item in the Supporting Information, Section S4). It also holds information on the thermostat, barostat, cutoff, restraints, and HMR, for which the MDDB input file has no structured field. What the record lacks, the force field and the water model, is not recorded in AMBER run files at all; it belongs to the system preparation, whose logs AmberMeta does not read.

**Table 4:** Items Asked for by a Reporting Checklist^10^ and by the MDDB Input File,^22^ the AMBER Files That Contain Them, and AmberMeta’s Output That Holds Them.

| item | AMBER source | AmberMeta output<sup>a</sup> |
|---|---|---|
| software and version<sup>b</sup> | mdout | *methods summary* |
| atom count | prmtop, mdout | *methods summary* |
| water molecules, ions | prmtop | *methods summary* (counts and atoms per water molecule; no concentration) |
| lipids, protonation states | prmtop | *methods summary* (residue names) |
| box dimensions | input coordinates | *methods summary* |
| ensemble<sup>b</sup> | mdin | *methods summary* |
| thermostat, barostat | mdin, mdout | *methods summary* (type and coupling) |
| target temperature<sup>b</sup> and pressure | mdin, mdout | *methods summary* |
| cutoff, electrostatics | mdin, mdout | *methods summary* |
| time step,<sup>b</sup> constraints, HMR | mdin, mdout, prmtop | *methods summary* |
| positional restraints | mdin | *methods summary* |
| frame interval<sup>b</sup> | mdin, mdout, trajectory | *methods summary* (write interval); *summary* (trajectory) |
| random seeds | mdin, mdout | *methods summary* |
| phases, runs, run lengths | *manifest*, mdout | *methods summary* |
| replicas and their simulated time | *manifest*, mdout | *methods summary* |
| force field,<sup>b</sup> water model<sup>b</sup> | not in run files | — |
| system preparation | not in run files | — |

<sup>a</sup> *Methods summary* (JSON and Markdown) written by plan; the *summary* holds the same items per run. Each setting in the *methods summary* carries its source: the mdin, the value echoed in the mdout, an AMBER default, or a value derived by AmberMeta. The full mapping is given in the Supporting Information (Section S4).
<sup>b</sup> Field of the MDDB input file.

Measured against the FAIR principles,^1^ the record provides rich metadata on how every run was performed (principles F2 and R1) and part of the provenance that R1.2 asks for: which file each run read and wrote, which program version ran it, and what was run as opposed to what was requested. Unlike forms filled in by hand, these metadata are checked against the output files of the runs. The *methods summary* is small enough, less than 20 kB, to be stored with the data or entered into the metadata of a repository, whereas the *summary* keeps every run for the archive.

For the record to be interoperable (principles I1 and I2), its terms must be linked to a shared vocabulary. AmberMeta’s interpreted settings use its own terms. MOLSIM, an ontology that our group is developing for biomolecular simulations,^37,38^ already defines classes for such settings, among them thermostats, barostats, ensembles, and the AMBER file formats, and it lists AMBER keywords such as ntt = 3 for some of them. The record of AmberMeta can thus serve as input for metadata annotated with MOLSIM terms; the annotation itself would be the task of an export (Discussion).

### Factual Basis of a Methods Section

The third use of AmberMeta is the Methods section. To test whether the *methods summary* can serve as its factual basis also when a language model writes the draft, three models (Claude Opus 5.5, Claude Sonnet 5.5, and Claude Haiku 4.5) each drafted the Methods section of the sample twice: once with a *plain request* for the Methods section and once with a *request with rules*, which forbid adding facts that the *methods summary* does not state and ask for placeholders and for the *findings* as questions to the authors. We checked every statement of the six drafts against the record and the run files (Methods).

With the explicit rules, the models added hardly anything of their own, but they did not always read the record correctly (Table 5). The largest model took the settings from the record, left placeholders for the force field, the ions, and the system preparation, and raised the time step of 4 fs on standard hydrogen masses as a question; its draft contained no error. The medium-sized model counted the two terminal caps as protein residues, gave the box at the start of production as the box after solvation, and stated a minimization that the record does not contain. The smallest model made no error but left out almost all placeholders and the 4 fs *finding*. Without the rules, the models also hallucinated content of their own, such as literature citations, the position of the ligand, or the identity of the protein read from a file name, and the smallest one read the oligosaccharide ligand chitohexaose as several ligands. Two of the misreadings concern values that the *methods summary* gives without saying what they exclude or when they were taken. The number of protein residues excludes the caps, and the box is the one at the start of production. Labeling both values in the *methods summary* would remove this source of error. Even so, a draft remains a draft: it has to be checked against the record, and what AMBER files do not contain has to come from the authors. The prompts, the drafts, and the check of every statement are given in the Supporting Information (Section S7).

**Table 5:** Statements in Methods Drafts Written by Three Language Models from AmberMeta’s *Methods Summary* of the Sample.

| request<sup>a</sup> | model<sup>b</sup> | unsup.<sup>c</sup> | contr.<sup>d</sup> | guess<sup>e</sup> | gaps<sup>f</sup> | 4 fs<sup>g</sup> |
|---|---|---|---|---|---|---|
| plain | Opus | 2 | 2 | 7 | 5 | yes |
| plain | Sonnet | 1 | 1 | **0** | 8 | yes |
| plain | Haiku | 3 | 3 | **0** | **0** | no |
| rules | Opus | **0** | **0** | 1 | 16 | yes |
| rules | Sonnet | 1 | 3 | **0** | 12 | yes |
| rules | Haiku | **0** | **0** | **0** | 1 | no |

<sup>a</sup> *Plain request*: a request for the Methods section. *Request with rules*: the same request with rules against adding facts. One draft per request and model, from the *methods summary* of version 1.3.0.
<sup>b</sup> Claude Opus 5.5, Claude Sonnet 5.5, and Claude Haiku 4.5.
<sup>c</sup> Statements given as fact that the model’s input does not contain; added literature citations count as one.
<sup>d</sup> Statements that contradict the record or the run files.
<sup>e</sup> Values inserted in brackets or flagged as guesses.
<sup>f</sup> Bracketed placeholders without a value.
<sup>g</sup> Whether the draft raises the time step of 4 fs on standard hydrogen masses.

## Discussion

We present AmberMeta, a tool that turns the files of a finished AMBER project into a checked protocol and writes a record of the runs. Its three uses rest on this record. As a sanity check, AmberMeta reported 146 of 152 errors introduced into in-house projects and showed unfinished runs, unequal replicas, and runs whose files were not collected with them in an in-house test collection; effectively, AmberMeta performs simulation run audits by providing a machine-readable history of the runs checked against their AMBER outputs. As for metadata, its record holds most of the setup items of a journal’s reporting checklist for MD simulations^10^ and part of the MDDB input file, checked against the output files of the runs. As the basis of a Methods section, its *methods summary* let language models draft text that kept largely to the recorded values when the request forbade adding facts, but only the draft of the largest model was free of errors and omissions.

Unlike single-run extractors such as GROMACS MetaDump,^23^ AmberMeta relates the runs of a project to each other and checks them against their output files. This is what reveals the simulated time of segmented simulations, missing and unfinished segments, runs that read another restart file than stated, and replicas that stopped early. Unlike iBIOMES Lite,^15^ it assumes no fixed order of stages, and where a directory layout does not identify the replicas, the user declares them. Because AmberMeta works after the simulations have finished, it complements workflow systems that record provenance during execution^27,28^ and also covers simulation runs started by hand-written job scripts.

AmberMeta prepares checked metadata on the simulation part of a dataset before the data are archived or deposited, and it thereby complements repositories and storage solutions for MD data.^7,36^ At the same time, it does not assign persistent identifiers (asked for in F1 of the FAIR principles^1^) or licenses (R1.1), its JSON and YAML files follow no public schema or controlled vocabulary (I1, I2), and it does not yet export to community formats (R1.3). It also records neither the system preparation nor hash sums of the input files, which the provenance records envisioned for MDDB include.^4^ An export of the record to the MDDB input file or to a Workflow Run RO-Crate, which packages files with their provenance,^39,40^ is a natural next step; the relation “run B started from the restart file that run A wrote” maps onto the PROV-O relations prov:used and prov:wasGeneratedBy,^41^ to which the terms of such crates are aligned.^39^ A public schema that the record could follow would have to hold such relations between runs. The BioSim metadata schema, for example, cannot: it describes a simulation by one stage of each kind, from setup to production.^42^ To the best of our knowledge, it has no place for replicas that share the preparation runs and then continue as separate chains of production segments (Figure 2A). A schema for such records must instead express a graph of runs, as the *manifest* does. Annotating the interpreted settings with MOLSIM terms and the files with EDAM formats^37,43^ would make the record interoperable down to single settings. Records of this kind could also feed the data management plans of projects, which are meant to become machine-actionable.^44^

The record also changes how a Methods section can be written. The *methods summary* holds the values that a Methods section reports about how the simulations were run, together with their sources, and it states what the run files do not contain. Authors can write the section from it, fill in a template, or let a language model draft it; pipelines in other fields already generate such text from recorded provenance.^45,46^ Our test suggests how to do this best: give the model the *methods summary* with rules that forbid adding facts, and check the draft against the record. Without the rules, the models added content of their own, as expected for text generated from data.^47^ With them, the models added hardly anything of their own but still misread parts of the record, and the smallest model left out the 4 fs *finding*. We note that the test is small, with one project, one draft per condition, and models from one provider, and drafts vary between runs. Because the *methods summary* carries the values that AMBER echoed, including those left at their defaults, it can also show where a Methods section written by hand differs from how the runs were performed.

Some further comments are in order. AmberMeta currently reads only AMBER files, but its model of *Simulation*, *Phase*, and *Step* is not specific to AMBER, and readers for other engines can be added to it. AMBER run files generally do not name the force field or the water model, which are combined per molecule type when the system is built,^13^ so the authors still have to add these items, and the *methods summary* lists them as missing. AmberMeta checks the bookkeeping of the runs, not their physics, and does not assess equilibration or convergence. Settings that neither the mdin sets nor the mdout echoes are filled in with the defaults of current AMBER versions and marked as defaults, because older versions do not always share them.

A second group of comments concerns what discover infers. Replicas are inferred only from directory layouts. Replicas in layouts that discover does not recognize are declared in the *manifest* or the editor. Until they are declared, the check of numbered series treats identically named series in different replica directories, such as prod_0001 to prod_0200 in each, as one series. A segment missing from one replica then leaves no gap in this series, because another replica has a segment with the same index. Three of the six misses in the error injection arose this way. Once the replicas are declared, each replica is checked against the combined range of all replicas, and the missing segment is reported. Parallel systems in separate directories (apo/01 beside holo/01) are not drafted as one *Simulation* if their runs record different atom counts; discover then names the systems that it found, so that each can be drafted on its own. Parallel systems with the same atom count are still grouped as one set of replicas, which the user then separates. Where an mdout records no usable input file, for example because of a truncated path, discover links the run to the run before it in its directory, and no check can test that link, because AMBER recorded nothing to compare it with. A run without an mdin is bound to a topology without knowing its time step, so that of a standard and an HMR topology with the same atom count, the draft chooses the standard one. The check of numbered series does not report runs missing at the end of a series in every replica or series numbered by a prefix (01_min, 02_heat). Replica exchange runs are flagged but not modeled.

In summary, AmberMeta reconstructs the protocol of a finished AMBER project from its files, checks it against the output files of every run, and writes a record of what was run as a project audit. We expect it to be most useful when a simulation campaign ends, before its results are analyzed and published and before its data are archived or deposited. Authors could include a brief audit summary with the supporting information and explain any discrepancies that affect the reported sampling. One can envision that Journals or MD data repositories encourage authors to use tools such as AmberMeta before submission to check whether the run files support the simulation details reported. AmberMeta can be installed with pip install ambermeta.

## Methods

### Sample and Modified Copies

The sample is distributed with AmberMeta (tests/data/amber/md_test_files). We ran discover, validate ––manifest, and plan with AmberMeta 1.3.0 on the unmodified sample and on copies into which we had introduced one error each (Table 3). The exact modifications, the commands, and their outputs are given in the Supporting Information (Section S5). The wall time of each command was measured five times after one unrecorded run.

### In-House Test Collection and Error Injection

All work on the in-house projects was done by a script package (kit) that ran on a workstation in our laboratory and never wrote into the projects; it read each project through a tree of symbolic links (Supporting Information, Section S6). For the survey, the kit ran discover, validate –– manifest, and plan on every project with simulation runs. For error injection, it took the eight projects for which discover had been fastest in the survey and introduced the kinds of errors listed in Table 3, which cover the everyday errors of the Introduction, at randomly chosen runs of copies of them. Errors in files were followed by a fresh discover on the modified copy; errors in the *manifest* were introduced into the *manifest* drafted for the unmodified project. An error counted as detected when a new *finding* of an expected kind referred to the run that carried it; new *findings* on other runs were counted as side effects. The researchers who ran the projects reviewed the *findings* of the survey.

### Run Time Measurements

The survey and the error injection ran on a workstation with an Intel Core i7-8700 and 32 GB of memory, which read the files from the network file system of our laboratory. The wall time and the peak memory of each command were measured per project, once with the files read from the network file system and then in three repetitions with the files cached. Synthetic projects of *N* segments were built from the sample by linking its files under new names, and each was run twice. The run started when the workstation was idle; during the survey and the benchmark, its load average rose to about 8 (Supporting Information, Section S6).

### Methods-Drafting Test

Each of the three models received one system prompt and either the *plain request* or the *request with rules*, followed by the *methods summary* of the sample (Supporting Information, Section S7). Each model was run once per request through the command line of Claude Code without access to tools, on 2026-10-07. We checked every statement of every draft against the record and the run files and counted statements given as fact that the input does not contain, statements that contradict the record or the run files, values inserted as guesses, and placeholders. The check was prepared and verified with AI assistance (Claude Code).

### Implementation and Tests

AmberMeta is written in Python (version 3.9 or newer). Its base installation needs only the standard library; NetCDF files are read with netCDF4 or SciPy, and the editor runs on a local web server. An automated test suite of the Python package, run on Python 3.9 and 3.12, and of the editor checks every change to the code.

### Use of AI Tools

Artificial intelligence (AI) tools were used in three ways. First, the AmberMeta code was written with the assistance of the AI coding tools Codex (OpenAI) from November 2025 to March 2026 and Claude Code (Anthropic) from January to November 2026. M.B. specified the behavior of the software, directed and reviewed the changes. Second, the manuscript and the Supporting Information were edited, proofread, and supplemented with AI assistance (Claude Code, Anthropic) in September and October 2026 from the authors’ notes, decisions, results, and checked sources, and they were revised by the authors. Each section of the Supporting Information states how AI was used for it. No images were generated by an AI model. Third, in the Methods-drafting test, the language models were the objects of the study, and the check of their drafts was prepared and verified with AI assistance (Methods-Drafting Test). The authors reviewed all content and take full responsibility for it.

## Data and Software Availability

AmberMeta is available at https://github.com/CPCLab/ambermeta and can be installed from the Python Package Index with pip install ambermeta. The version described here is 1.4.0, archived at https://researchdata.hhu.de/handle/entry/245. The results were obtained with version 1.3.0. Version 1.4.0 adds the acknowledgement of checked *findings* with a note and a Markdown version of the *methods summary*, no longer drafts systems with different atom counts as one *Simulation*, and words several messages more plainly. It is currently distributed under the Business Source License 1.1, which permits non-production use, and will be released under the MIT License upon publication in a peer-reviewed journal. The sample data used in the Results are included in the repository (tests/data/amber/md_test_files). The in-house projects belong to a collection that members of our laboratory assembled to develop and test tools for research data management. They are partly unpublished and not publicly available; the procedures and anonymized results per project are given in the Supporting Information.

## Supporting Information Available

How AmberMeta reads AMBER files (S1), drafts the protocol (S2), and checks it (S3), with flowcharts of its rules; the record and its coverage of reporting items (S4); the sample project (S5); the procedure and results on the in-house test collection (S6); and the Methods-drafting test (S7) (PDF). Data files: command outputs and modified copies of the sample; runs, run times, *findings*, and error injection per in-house project; and the input, prompts, drafts, and statement check of the Methods-drafting test (ZIP).

## Author Information

ORCID: Michele Bonus, Rocco Gentile, Alexander Schug, Holger Gohlke.

### Author Contributions

M.B.: Conceptualization, Methodology, Software, Validation, Formal analysis, Investigation, Visualization, Writing – original draft, Writing – review & editing; R.G.: Data curation, Investigation, Validation, Writing – review & editing; A.S.: Resources, Validation, Funding acquisition, Writing – review & editing; H.G.: Conceptualization, Validation, Resources, Supervision, Project administration, Funding acquisition, Writing – review & editing. All authors have approved the final version of the manuscript.

## Acknowledgement

We are grateful to the members of our labs at Heinrich Heine University Düsseldorf and Forschungszentrum Jülich for providing the MD simulation files of the in-house projects. The work was supported by Helmholtz Metadata Collaboration Project Funds (ELN-RO (ZT-I-PF-3-091)), Helmholtz Gateway Funds, and Helmholtz HFMI Project Funds (PROFOUND). The use of AI tools is described in the Methods (Use of AI Tools).

---

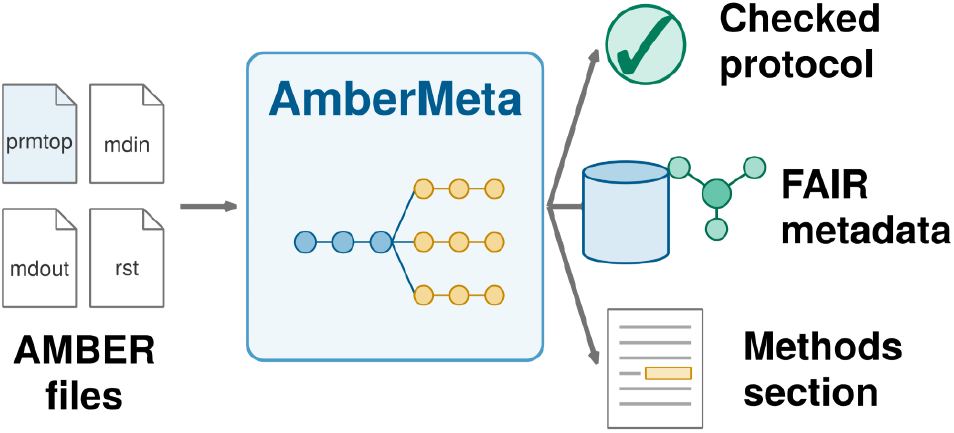
TOC Graphic.

