## Supporting Information for "AmberMeta: From AMBER Simulation Files to Checked Protocols and FAIR Metadata"

**About this Supporting Information.** Sections S1 to S4 explain how AmberMeta works: what it reads from AMBER files, how it drafts a protocol, what it checks, and what it writes. These actions are governed by rules. The sections give each rule with the reason for it; the complete rules are part of the documentation that comes with the software. The AMBER Reference Manual defines the file formats and settings referred to here.[S1]

Sections S5 to S7 describe the evaluation: the sample project, the in-house test collection, and the Methods-drafting test. Command outputs, values per project, run times, and the drafts of Methods sections written by the language models are provided as data files (Table S1); the sections of this Supporting Information concentrate on what these files show.

Each section states how AI tools were used for it. The terms *Simulation*, *Phase*, *Step*, *manifest*, *finding*, *summary*, and *methods summary* are defined in Table 2 of the main text and set in italics.

The in-house projects belong to a test collection that members of our laboratory assembled to develop and test tools for research data management. The projects belong to the researchers who ran them and are partly unpublished; they are therefore referred to by anonymized identifiers (`sys001` to `sys050`), and no file, directory, or system names from them are given. The exceptions are `sys021` and `sys022`, the published simulations of the human taurine transporter TauT and its variant p.A78E.[S2]

**Table S1:** Data Files of the Supporting Information (`si_files.zip`)

| file | content | section |
|---|---|---|
| `S5_sample/commands_and_outputs.txt` | commands run on the sample and their outputs | S5 |
| `S5_sample/modified_copies.csv`, `..._outputs.txt` | the modified copies of the sample, their *findings*, and the outputs | S5 |
| `S5_sample/segment_statistics.csv`, `timing.csv` | statistics per segment; wall time per command | S5 |
| `S6_inhouse/projects.csv` | runs, replicas, files, size, simulated time, run time, and memory per project | S6 |
| `S6_inhouse/survey_findings.csv` | *findings* per project and kind | S6 |
| `S6_inhouse/injection_scenarios.csv`, `injection_results.csv`, `injection_misses.txt` | scenarios, results per project and scenario, misses and side effects | S6 |
| `S6_inhouse/synthetic_scaling.csv`, `workstation.txt` | run time of synthetic projects; hardware and software | S6 |
| `S7_methods_drafting/methods_summary_sample.json`, `prompts.txt` | input of the language models | S7 |
| `S7_methods_drafting/drafts/*.md`, `statement_check.csv`, `summary.csv` | the six drafts; the check of every statement | S7 |

### S1 Reading AMBER Files

AI use: Proofread and supplemented with AI assistance (Claude Code) using the source code.

To describe a run as a Methods section would, AmberMeta needs to know which file belongs to which run and what each file says about that run. This section explains both.

#### S1.1 File Types

AmberMeta first assigns a type to every file (Figure S1). Most extensions are unambiguous, but two cases need care. (i) The extension `.out` is also used by job schedulers and other programs, so an `.out` file counts as an AMBER output only when its beginning contains a text that AMBER writes at the top of every output, such as the program banner or the File Assignments block; job logs named `slurm-*.out` are never read as outputs. (ii) NetCDF restart files and trajectories have the same file format, and AmberMeta tells them apart by their extensions alone. A file that is missing or cannot be opened is noted on its run, and AmberMeta continues with the remaining files.

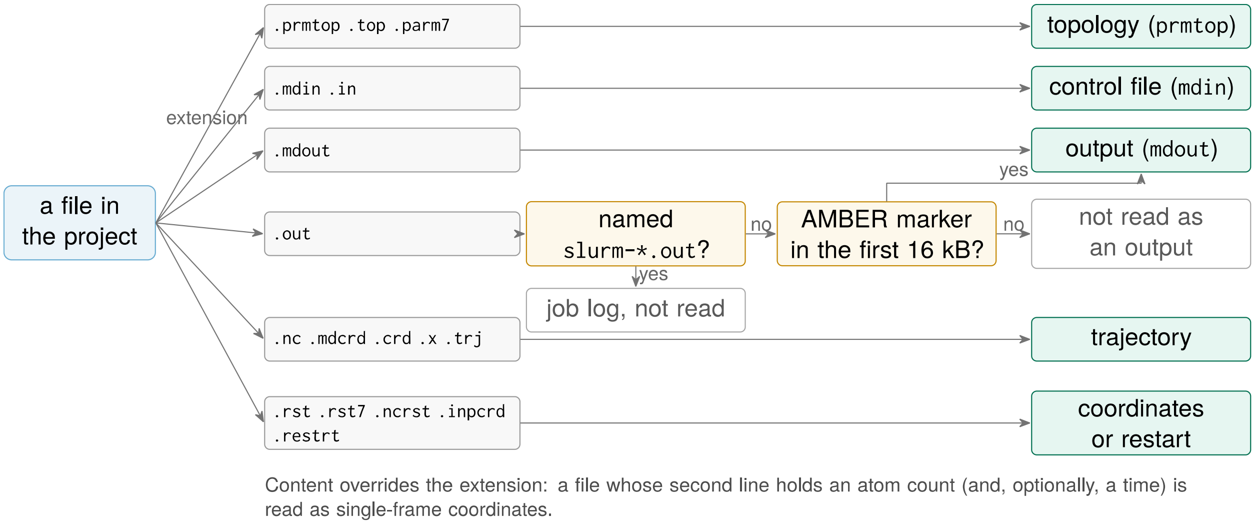


**Figure S1:** How AmberMeta recognizes the type of a file. The extension decides for most files, but two cases need care. An `.out` file may also be a job log, so it counts as an AMBER output only when its beginning contains a text that AMBER writes. NetCDF restart files and trajectories have the same file format and are told apart by their extensions alone. In addition, for single-frame coordinate files, the content overrides the extension (note at the bottom).

#### S1.2 Contents of Each File

Each type of file contributes one part of the description of a run (Table S2).

**Table S2:** What AmberMeta Reads from Each Type of File, and What It Is Used For

| file | what AmberMeta reads | used for |
|---|---|---|
| topology (`prmtop`) | atom and residue counts, residue names, charges, masses, the box written when the system was built | the system (residues by class, water molecules and ions, net charge); the hydrogen mass repartitioning (HMR) label |
| control file (`mdin`) | every namelist setting, kept as written | interpreted settings (Table S3); methods that need separate reporting |
| output (`mdout`), header | the files the run read and wrote (File Assignments); the settings as AMBER used them; the random seed | links between runs; comparison with the `mdin`; defaults; random seeds |
| output (`mdout`), body | every energy record (time, temperature, pressure, density, energy); the completion marker | simulated time; statistics per run; finished or not |
| restart files, NetCDF trajectories | simulation time, box, atom count | continuity in time; atom counts |

**Hydrogen masses.** Whether a time step of 4 fs is safe depends on the hydrogen masses, so AmberMeta labels every topology as standard or HMR. A topology counts as HMR when its heaviest hydrogen weighs at least 2.0 amu, or at least 1.5 amu while other hydrogen atoms kept a mass near 1.008 amu. Hydrogen atoms are identified by their atomic number or, where a topology lacks it, by their name and mass.

**Requested and used settings.** The `mdin` states what was requested, and the header of the `mdout` what AMBER used, including the values that AMBER filled in for settings that the `mdin` left out. AmberMeta reads both: their comparison reveals an `mdin` that was edited after its run, and the `mdout` supplies the defaults that a Methods section has to report. Where neither file states a setting, AmberMeta uses the default of current AMBER versions (Table S3), and the *methods summary* marks it as a default. The random seed is taken from the `mdout`, because many `mdin` files request a random seed (`ig = -1`) and only the `mdout` records the value that was drawn. Flags mark methods that need separate reporting: replica exchange, Gaussian accelerated MD, free energy calculations, constant pH, constant redox potential, QM/MM, and NMR-type restraints. AmberMeta records them but does not model them explicitly; a replica exchange run, for example, is one *Step* like any other.

**Table S3:** Interpretation of the Main `&cntrl` Settings and the Values Used When a Setting Is Absent

| setting | interpretation | when absent |
|---|---|---|
| `imin` | 0, dynamics; 1, minimization | 0 |
| `nstlim`, `dt` | number of integration steps; time step | 1; 0.001 ps |
| `ntt`, `temp0` | thermostat (0, none; 1, Berendsen; 2, Andersen; 3, Langevin; 11, Bussi; and others); target temperature | 0; 300 K |
| `ntp`, `barostat` | pressure coupling (isotropic, anisotropic, semi-isotropic); barostat (1, Berendsen; 2, Monte Carlo) | 0; 1 |
| `ntb`, `igb` | periodic boundaries; implicit solvent | from `igb` and `ntp`; 0 |
| `ntc`, `cut` | bond constraints; nonbonded cutoff | 1; 8 Å (9,999 Å with `igb` > 0) |
| `ntr`, `restraintmask`, `restraint_wt` | positional restraints, their atoms, and their force constant | 0 |
| `ntpr`, `ntwx`, `ntwr` | intervals of energy records, trajectory frames, and restart files | 50; 0; `nstlim` |
| ensemble | derived from `ntb`, `ntt`, `ntp`, and `igb` (NVE, NVT, NPT, NPH) | follows |

**Simulated time and statistics.** The simulated time of a run is the time of its last energy record minus the time at which the run started. AmberMeta counts what was run, not what the `mdin` requested; runs without output, minimizations, and queued runs contribute none. The mean and standard deviation of temperature, pressure, density, and energy are accumulated over all energy records with Welford's algorithm,[S3] which needs constant memory for an output of any length. A run counts as finished when its `mdout` contains the timing summary that AMBER writes at the end of a run.

**Trajectories.** To check continuity, AmberMeta needs the times of the first and the last frame of a trajectory, not its coordinates. Because reading every frame of a NetCDF file is slow on network file systems, it reads the times of only five frames: the first, the last, and the three at one quarter, one half, and three quarters of the trajectory. It reads all frames only when these five do not lie on one constant frame interval, as in a trajectory that was cut off. Reading NetCDF files requires the netCDF4 or SciPy package.

### S2 Drafting the Protocol with `discover`

AI use: Proofread and supplemented with AI assistance (Claude Code) using the source code.

The `discover` command drafts the *manifest* so that the user does not have to write it, and its rules decide what the user has to check. This section gives the rules in the order in which they act: grouping files into runs, assigning roles, linking runs, and recognizing replicas.

#### S2.1 Runs and Roles

Files with the same path apart from the extension form one run, for example `md_0002.mdin`, `md_0002.mdout`, `md_0002.nc`, and `md_0002.rst`; a group with an `mdin` or an `mdout` becomes a *Step*. A *Step* with an `mdin` but no `mdout` was prepared and did not run, and it is marked as queued. From version 1.4.0, runs whose `mdout` files record different atom counts are not drafted as one *Simulation*: `discover` names the systems that it found, each by the shortest directory that holds its runs, so that the user can draft each on its own or ask for one *Simulation* explicitly (option `--as-one-simulation`). Runs are taken in natural order, in which `prod_2` precedes `prod_10`.

Each run gets its role from the strongest evidence available (Table S4). An `mdin` that requests a minimization is unambiguous and decides first. Otherwise, words in the directory and file names decide, from the top directory down. They are matched only as whole words, so that `prod1` and `eq0001` carry their cue, whereas a directory named `minor` does not count as `min`. Without such a word, the settings decide. Consecutive runs with the same role form a *Phase*. Because names follow no common rule, the role is the first thing that the user should check in the draft.

**Table S4:** Evidence for the Role of a Run, in the Order in Which It Is Used

| evidence | role | examples and remarks |
| --- | --- | --- |
| 1. the `mdin` requests a minimization (`imin = 1`) | minimization | decides regardless of the name |
| 2. a word in the directory or file name | | first matching directory from the top, then the file name |
| `min`, `minim`, `minimization`, `em` | minimization | `01_min`, `em/` |
| `heat`, `warm`, `therm`, `anneal` | heating | `heat_eq` counts as heating |
| `eq`, `equi`, `equil`, `equilibration` | equilibration | `eq0001`, `equil/` |
| `prod`, `production` | production | `prod1`, `nvt_prod_0001` |
| `nvt`, `npt` (weaker) | equilibration | used only without the cues above: `npt_02` |
| 3. the settings | | without any word in the name |
| positional restraints (`ntr = 1`) | equilibration | |
| start at 50 K or below, `temp0` higher | heating | |
| more than 500,000 integration steps | production | |

Words such as `md` and `run` are not cues, because they are used for every stage.

#### S2.2 Links Between Runs

A Methods section describes the order of the stages; AmberMeta also needs to know which run continued from which restart file. File names are an unreliable guide to this. In the first runs of AmberMeta on the in-house collection, linking by file order put runs out of the order in which they ran, for example where directories sorted by name placed equilibration before minimization. `discover` therefore links each run to the run whose restart file AMBER recorded as its input (Figure S2). The recorded path often belongs to the computer on which the run was executed; AmberMeta then looks for restart files of the same name and, among several, chooses the one in the run's own directory or replica. Only where AMBER recorded no usable input, for example because a long path was cut off, is a run linked to the previous run in its directory.

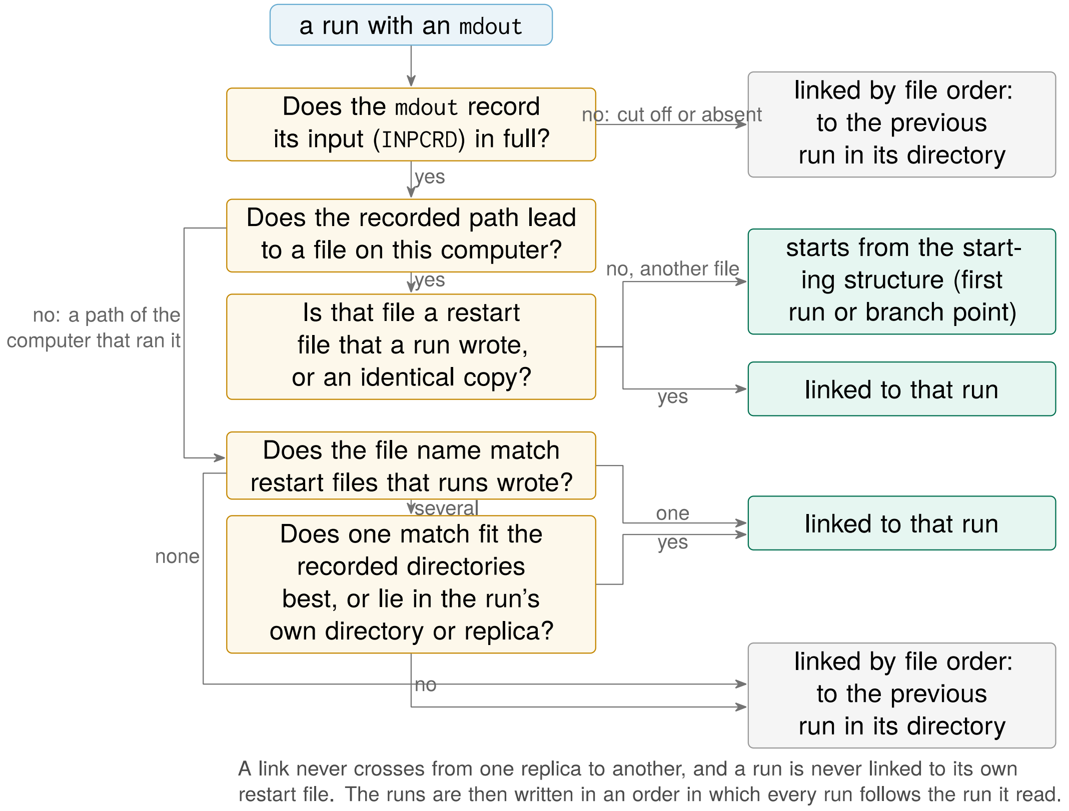


**Figure S2:** How `discover` links a run to the run whose restart file it read. The input that AMBER recorded in the `mdout` decides; file order is the fallback.

The starting structure is the coordinate file that the first run records as its input; where it records none, it is the first single-frame coordinate file that no run wrote. Identical copies of one file, such as a starting restart file copied into every replica directory, count as one file.

#### S2.3 Replicas

`discover` recognizes replicas from the directory layout (Figure S3): directories at the same level that hold the same series of runs become replicas, named after the directory, and directories with the shared preparation stay outside them. Layouts that do not identify the replicas unambiguously are left to the user, because a wrong guess would compare runs that do not belong together. Among them are replicas whose directories are named differently in different stages, such as production directories `run1` to `run4` beside `equilibration/equi1` to `equi4`, and replicas grouped by a condition, such as `300K/rep1` beside `310K/rep1`. For new projects, the documentation recommends a layout that `discover` recognizes: one directory per stage, the replicas as subdirectories with the same names in every stage (`equilibration/run1`, `production/run1`), and the replica named only in the directory, not in the file names. In the *manifest*, the replica of a run is stored as its `lineage` tag.

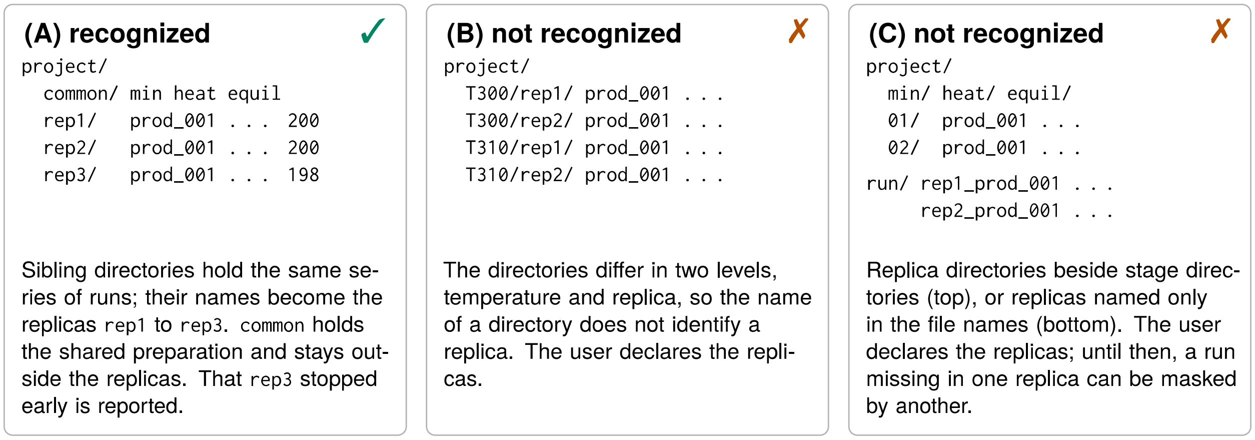


**Figure S3:** Directory layouts from which `discover` does (A) or does not (B, C) infer replicas. In B and C, the user declares the replicas in the editor or the *manifest*.

##### S2.4 Output of `discover`

`discover` prints the draft and a list of suggestions. Suggestions marked `applied` were carried out and should be checked: the starting structure, the roles, and the replicas. Suggestions marked `needs_you` are questions that only the user can answer: whether a topology is an HMR topology, which runs are replicas in an ambiguous layout, and whether indices missing from a numbered series are real gaps. The output for the sample is given in the data files (Section S5).

#### S3 Checks of `validate`

AI use: drafted with AI assistance (Claude Code) from the source code and the authors' design decisions.

The `validate` command checks the protocol against what AMBER recorded. Each check targets an error that occurs in everyday work and that the control files do not show (Table S5).

**Table S5:** The Checks of `validate`, What They Compare, and the Errors They Catch

| check | compares | catches | *finding* (kind) |
|---|---|---|---|
| recorded input | the input that the *manifest* declares with the input that the `mdout` records, file and directory | a run that read another restart file than the protocol states | `input_mismatch` |
| completion | the `mdout` with the timing summary that AMBER writes at the end | a run that stopped early or is still running | `unfinished_run` |
| atom counts | topology, coordinates, `mdout`, trajectory | a run with the topology of another system | `step_check` |
| `mdin` and `mdout` | steps, time step, run length, frame interval | an `mdin` edited after its run | `step_check` |
| trajectory span | the time span of the trajectory with the length of the run | a truncated or foreign trajectory | `step_check` |
| time step and hydrogen masses | a time step above 2 fs with the label of the topology | a missing HMR topology or an unsupported time step | `step_check` |
| continuity | the start of a run with the end of the run it continues | an overwritten restart file (overlap); missing simulation time (gap) | `continuity_gap` |
| branches | runs of one replica that continue from the same restart file | a job resubmitted with an older restart file | `continuity_gap` |
| numbered series | the indices of a series within and across replicas | a missing segment; a replica that stopped early | `missing_run` |
| replicas | atom counts, minimization only, per role the time step, cutoff, thermostat, and pressure coupling, and the target temperature in production | replicas that do not describe the same system under the same conditions | error or warning |
| random seeds | the seeds of runs that continue from one restart file | replicas that are not independent | warning |
| files | every file that the *manifest* names | missing or unreadable files | error |

Two checks need more than a comparison of values (Figure S4). (i) For the recorded input, a comparison of file names is not enough, because replicas name their restart files alike, and AmberMeta also compares the directory. (ii) For numbered series, the replicas of a series are compared over their combined range, so that a replica that stopped before the others is reported; runs missing at the end of every replica are not, because nothing shows that they should exist.

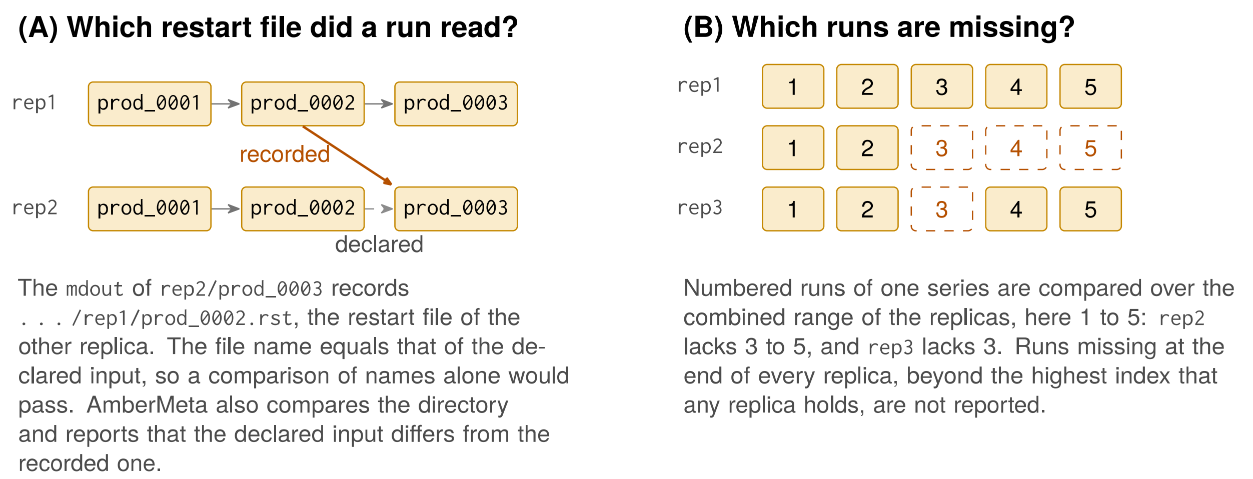


**Figure S4:** Two checks on small examples: (A) the recorded input across replicas and (B) numbered series across replicas.

Small differences are tolerated where the files cannot be exact (Table S6). Continuity accepts differences of up to 0.1 ps or half a frame interval, because NetCDF trajectories store times in single precision and the last frame of a run need not coincide with its end. Gaps that the user declares in the *manifest* are accepted, whereas overlaps are always reported. Replicas are compared role by role, and the target temperature only in production. Some protocols equilibrate each replica at a slightly different temperature, such as 299.9 K, 300.0 K, and 300.1 K, before all continue at a common production temperature; such replicas are not reported. This differs from replica exchange, in which replicas at different temperatures exchange configurations during the run; AmberMeta flags replica exchange runs but does not model them.

**Table S6:** Thresholds and Tolerances in AmberMeta 1.3.0

| quantity | rule |
|---|---|
| HMR label | heaviest hydrogen ≥ 2.0 amu, or heaviest ≥ 1.5 amu and lightest ≤ 1.1 amu |
| time step that needs HMR | above 2 fs |
| continuity | difference within $\max(0.1\,\text{ps}, 0.5\,\Delta t_{\text{frame}})$, or within a declared window |
| trajectory span | within $\max(0.001\,\text{ps}, \Delta t)$ of the span that `nstlim`, `dt`, and `ntwx` predict |
| numbered series | at least two runs of one series; indices between the lowest and the highest present in the replicas of the series |
| `.out` file read as `mdout` | AMBER marker within the first 16,384 bytes |

For use in scripts, `validate` ends with a nonzero exit code when files named in the *manifest* are missing or cannot be opened, when atom counts differ within or between replicas, or when only some of the replicas ran dynamics. From version 1.4.0, a *finding* that the user acknowledged with a note in the *manifest* is listed as acknowledged and does not count for the strict option. With the option `--strict`, every *finding* also leads to a nonzero exit code, so that a script can stop before a project is archived. The `plan` command writes its output in every case, even when it ends with a nonzero exit code.

### S4 The Record

AI use: Proofread with AI assistance (Claude Code).

The `plan` command writes three files, each for one use. The *summary* (JSON or YAML) is the full record for the archive: for every run, its role, replica, files, parsed settings, the values that its `mdout` echoes, its simulated time, and its *findings*, with totals per replica, per role, and overall.

The *methods summary* (JSON; from version 1.4.0 also as Markdown) condenses the *summary* into what a Methods section reports. Its parts describe the software, the system, the replicas, the protocol, the continuity, the *findings*, and the items that AMBER run files do not contain. The protocol lists the *Phases* in the order in which they ran, and each setting is stated once per *Phase* with the number of runs that share it, the values of the runs that differ, and its source: `mdin`, `mdout` (echoed by AMBER because the `mdin` left it at its default), `default` (stated by neither file), or `derived` (computed by AmberMeta, such as the ensemble). The *methods summary* of the sample is given as a data file (Section S7). The table of run statistics (CSV) has one row per run with its simulated time and the mean and standard deviation of temperature, pressure, density, and total energy.

#### S4.1 Coverage of Reporting Items

Tables S7 to S9 compare the record with a reporting checklist for MD simulations[S4] and with the MDDB input file.[S5] Many of these items correspond to classes of the MOLSIM ontology,[S6] which could annotate them in a future export (main text, Discussion).

**Table S7:** Items of a Reporting Checklist for MD Simulations and Their Coverage by AmberMeta's Record: Items 1 to 3 (AmberMeta 1.3.0, Sample Project)

| item[a] | element asked for | AmberMeta output (field) | AMBER source | cov.[b] |
|---|---|---|---|---|
| 1a | evidence that the measured property equilibrated | observed temperature, pressure, and density per phase (methods summary); mean and standard deviation per run (statistics table); no time course | `mdout` | P |
| 1b | split into equilibration and production | phases in execution order with their roles, runs, and simulated time (methods summary) | manifest, `mdout` | C |
| 1b | amount of production data analyzed | not covered; simulated time per phase and replica (methods summary) | `mdout` | N |
| 1c | at least three simulations per condition | number of replicas and simulated time per replica (methods summary); no statistics | directory layout or manifest | P |
| 1d | independence from the initial configuration | random seeds per phase and the run from which each replica continues (methods summary) | `mdin`, `mdout`, manifest | P |
| 2a | connection to experiments | not covered | — | N |
| 3a | system with membranes, nucleic acids, glycans, … | residues by class: protein, nucleic acid, lipid, water, ion, other (methods summary) | `prmtop` | P |
| 3b | force field and water model | not covered; atoms per water molecule and force-field hints only (methods summary) | not in run files | N |
| 3b | implicit or explicit solvent | solvent type, periodic boundaries (methods summary) | `prmtop`, `mdin` | C |
| 3c | enhanced-sampling parameters | special methods per phase (methods summary); verbatim settings and further namelists (summary) | `mdin` | P |

[a] Numbering of the checklist.[S4]

[b] C, covered; P, partly covered or only in the summary; N, not covered.

**Table S8:** Items of a Reporting Checklist for MD Simulations and Their Coverage by AmberMeta's Record: Item 4 (AmberMeta 1.3.0, Sample Project)

| item[a] | element asked for | AmberMeta output (field) | AMBER source | cov.[b] |
|---|---|---|---|---|
| 4a | box dimensions | box of the input coordinates (methods summary) | input coordinates, `mdout`, `prmtop` | C |
| 4a | total number of atoms | atom count of each topology (methods summary); atom counts per file (summary) | `prmtop`, coordinates, `mdout`, trajectory | C |
| 4a | number of water molecules | water molecules and atoms per molecule (methods summary) | `prmtop` | C |
| 4a | salt concentration | ion counts only; no concentration, counter-ions not separated (methods summary) | `prmtop` | P |
| 4a | lipid composition | lipid residue names with counts (methods summary) | `prmtop` | P |
| 4b | protonation states | residue names only (methods summary) | `prmtop` | P |
| 4b | structural restraints | mask and force constant per phase, with the series of force constants over runs (methods summary) | `mdin` | C |
| 4b | nonbonded cutoff | cutoff with its source (methods summary) | `mdin`, `mdout` | C |
| 4b | thermostat | type, coupling constant, and target temperature with their sources (methods summary) | `mdin`, `mdout` | C |
| 4b | barostat | type, scaling, target pressure, relaxation time, and compressibility with their sources (methods summary) | `mdin`, `mdout` | C |
| 4b | time step, constraints, ensemble, run length, HMR | time step, constraints, ensemble, requested run length, HMR label (methods summary) | `mdout`, `mdin`, `prmtop` | C |
| 4b | long-range electrostatics | method as echoed in the `mdout` (methods summary); grid parameters not read | `mdout` | P |
| 4c | simulation software and version | program and version, GPU models, run dates (methods summary) | `mdout` | C |
| 4c | analysis software and version | not covered | — | N |
| 4d | input files and initial and final coordinates provided | not covered by the outputs; the manifest names the files of every run, without hash sums | manifest | N |
| 4e | custom code and parameters provided | not covered | — | N |

[a] Numbering of the checklist.[S4]
[b] C, covered; P, partly covered or only in the summary; N, not covered.

**Table S9:** Fields of the MDDB Input File and Their Coverage by AmberMeta's Record

| MDDB field[a] | AmberMeta output (field) | cov.[b] |
|---|---|---|
| `program`, `version` | program and version from the `mdout` (methods summary) | C |
| `timestep` | time step (methods summary) | C |
| `framestep` | write interval times time step (methods summary); frame interval of the trajectory (summary) | C |
| `ensemble` | ensemble (methods summary; own vocabulary) | C |
| `temp` | target temperature with its source (methods summary) | C |
| `method` | special methods per phase (methods summary) | P |
| `wat` | atoms per water molecule only (methods summary) | P |
| `type`, `ff` | not covered | N |
| input topology, structure, and trajectory paths | topologies, starting structure, and trajectory of every run (manifest) | C |
| `mds` (one entry per MD) | replicas with their simulated time and branching (methods summary) | P |
| administrative, reference, annotation, and viewer fields (33) | not covered | N |

[a] Top-level keys of the template of the MDDB input file (repository `mmb-irb/MDDB-workflow`, commit 91f39de, accessed 2026-10-02).[S5]

[b] C, covered; P, partly covered; N, not covered. The MDDB input file has no field for the thermostat, barostat, cutoff, restraints, HMR, or hardware, which AmberMeta records; they fit only its free-form field `metadditions`.

### S5 The Sample Project

AI use: Proofread and checked with AI assistance (Claude Code) from the outputs of AmberMeta, which are given unchanged in the data files.

This section makes the example of the main text reproducible. The sample is distributed with AmberMeta (`tests/data/amber/md_test_files`) and consists of 18 files (34 MB): the topology, the coordinate file written by `tleap`, the restart file from which the first segment started, and the `mdin`, `mdout`, and restart file of each of the five segments; the trajectories are not part of the sample. The segments were run with `pmemd.cuda` from AMBER 22[S7] for 5,000,000 steps of 4 fs each, with a Langevin thermostat at 300 K, isotropic pressure coupling, bonds to hydrogen constrained, and a cutoff of 9 Å. The system of 64,528 atoms contains 14,659 water molecules, 34 potassium ions, and 38 chloride ions.

#### S5.1 Procedure

Every case was run on a fresh copy of the 18 files with AmberMeta 1.3.0, on the workstation and by the script package described in Section S6: `discover` wrote the draft, `validate` checked it with and without the strict option, and `plan` wrote the record. For the modified copies, one error of Table 3 of the main text was introduced into each copy. The commands, their outputs, and the exact modifications are given in the data files; recorded paths of the cluster on which the runs were executed are shortened there to `.../` followed by the file name.

#### S5.2 Results

`discover` drafted the protocol as the runs were executed: one production *Phase* of five segments, each continuing from the restart file of its predecessor. The record gives 100 ns of simulated time, and the means of temperature and density per segment agree within their standard deviations (Figure S5). The one *finding*, the time step of 4 fs on standard hydrogen masses, is discussed in the main text. In each modified copy, AmberMeta reported the introduced error by the *finding* expected for it, and the strict check failed (`modified_copies.csv`). Each command took less than a second.

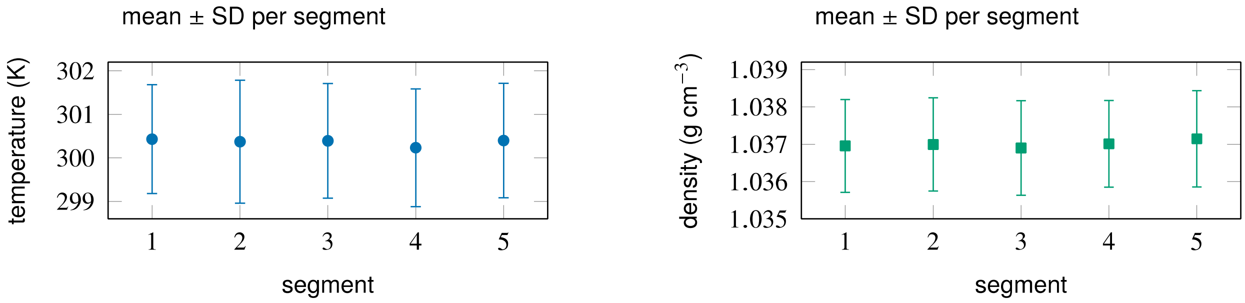


**Figure S5:** Temperature and density per segment of the sample (mean and standard deviation over the energy records of each segment), from the table of run statistics.

### S6 The In-House Test Collection

AI use: The scenarios of the error injection were realized with AI assistance from the author's instructions to cover the errors of Table 3 of the main text.

This section describes how AmberMeta was applied to the in-house test collection and summarizes what the data files give per project.

#### S6.1 Procedure

All work on the collection was done by a script package (kit) that ran on a workstation in our laboratory and read the projects from the network file system on which they are stored (`workstation.txt`). The kit never wrote into the projects: it read each project through a tree of symbolic links, and a modified file replaced its link with a modified copy. A listing of all files before and after the run showed that none was added, removed, or changed. The kit ran three stages.

**Survey.** For each project with simulation runs, the kit ran `discover`, `validate`, and `plan`, as a data steward would before archiving the project. The researchers who ran the projects reviewed the resulting *findings*.

**Error injection.** The kit took the eight projects for which `discover` had been fastest in the survey. For each kind of error (`injection_scenarios.csv`), it listed every place in a project where the error could occur, mostly chains of three runs in one directory of which each continues from the previous one, and drew up to three places per project at random, with a generator seeded with the project and the kind of error, which makes the draw repeatable. After an error was introduced into a file, `discover` drafted a new *manifest* from the modified copy. Errors in the *manifest* were introduced into the *manifest* drafted for the unmodified project. An error counted as reported when a new *finding* of the expected kind referred to the run that carried it. New *findings* on other runs were counted as side effects.

**Run time.** The kit measured the wall time and peak memory of each command per project, first with the files read from the network file system and then in three repetitions with the files cached. To measure the cost per run independently of the projects, it also built synthetic projects of increasing size by linking the files of the sample under new names. Each command ran four times per project: the first run read the files from the network file system, and the median of the other three gives the time with the files cached. The run started when the workstation was idle; during the survey and the benchmark, its load average rose to about 8 (`workstation.txt`).

#### S6.2 Results

**Error injection.** Of 156 drawn variants, 4 could not be made (for example, an `mdin` without `nstlim`), and AmberMeta reported 146 of the 152 introduced errors (`injection_results.csv`). The six misses had three causes (`injection_misses.txt`). (i) Three occurred in a project whose replicas have one directory for production at the top level and a differently named one for equilibration inside a common equilibration directory, a layout from which AmberMeta infers no replicas (Figure S3C); a run with the same index in another replica then masked the gap. (ii) Two occurred where the next run had been set up but had not yet run. Only a run that has run records which file it read, so nothing showed the error; in one of the two, the modified run was also the last of its numbered series (Figure S4B). (iii) One concerned a restart file overwritten in a chain of minimizations; minimizations do not advance the simulation time, so there was no time to compare. Where an error created a branch, AmberMeta reported both runs of the branch, as intended. Side effects were rare: four variants, all in one project, gave one or two further *findings* on neighboring runs.

**Survey.** `discover` read all projects without a failed file. The *findings* fall into these groups, given per project in `survey_findings.csv`: runs that had not finished; replicas of different length, which the files cannot tell apart from replicas that were meant to differ; runs at 4 fs on a standard topology, either with no HMR topology in the collection or because the draft bound runs without `mdin` files to the standard one of two topologies with the same atom count; runs whose producing runs were not collected; trajectories that do not cover the length of their run; replicas that ran production at different target temperatures; a time step in an `mdin` that differs from the one its `mdout` records, presumably because the `mdin` was edited after the run; and an input file of the system-building program that was taken as a run. Ten of the 19 projects had no *finding*.

**Run time.** The run time grows linearly with the number of runs and is dominated by reading files from the network file system (`projects.csv`, `synthetic_scaling.csv`; Figure 4C of the main text). The peak memory is set by the size of the topology file.

### S7 The Methods-Drafting Test

AI use: the language models were the objects of this test. Drafts were realized by the large language models and reviewed by the authors. Draft content was further cross-checked against actual simulation data with AI assistance (Claude Code).

This section describes how we tested whether the *methods summary* can serve as the factual basis of a Methods section when a language model writes the draft.

#### S7.1 Input and Requests

The input was the *methods summary* of the sample (`methods_summary_sample.json`). Both requests used the same system prompt, which describes the model as an assistant to researchers in MD simulation who write a journal article (`prompts.txt`). The *plain request* asks for the Materials and Methods section and appends the *methods summary*. The *request with rules* asks for the part of the section that describes how the simulations were run and sets four rules: use only information that the *methods summary* states; write a bracketed placeholder for every item that a Methods section would state but the *methods summary* does not contain; list the *findings* as open questions after the text; and write in the past tense, in plain prose.

#### S7.2 Procedure and Check

Claude Opus 5.5, Claude Sonnet 5.5, and Claude Haiku 4.5 each received both requests once, through the command line of Claude Code without access to tools, on 2026-10-07. We split every draft into its statements and checked each against the *methods summary* and the run files (`statement_check.csv`). A statement was supported when the input states it, unsupported when it is given as fact but the input does not contain it, contradicting when it disagrees with the record or the run files, a guess when a value was inserted in brackets or marked as uncertain, and a placeholder when it left a value to the authors. Table 5 of the main text gives the counts per draft (`summary.csv`).

#### S7.3 Results

With the *request with rules*, the drafts took the settings from the record and added hardly anything of their own. The errors were misreadings of the record, all in the draft of the medium-sized model: it counted the N-terminal acetyl and C-terminal N-methyl caps among the 362 protein residues, which the record counts without them, gave the box at the start of production as the box after solvation, listed seven residues as six, and stated a minimization that the record does not contain. The smallest model left out almost all placeholders and did not raise the time step of 4 fs on standard hydrogen masses.

With the *plain request*, the models added content of their own: literature citations, the position of the ligand in the protein, the identity of the protein read from a file name and stated as fact, and a reason for a setting that the record does not give; one draft also credited AmberMeta with suggesting a force field and a water model, which the record does not name. The same misreadings of the residue count and the box occurred there, and the smallest model also read the oligosaccharide, one molecule of seven residues, as several ligands. Both kinds of misreading, the residue count and the box, concern values that the *methods summary* gives without saying what they exclude or when they were taken; labeling both values would remove this source of error. The six drafts are given as data files (`drafts/`). The check was prepared with AI assistance and verified in a second, independent pass; statements that misread the record count as contradictions.
